# Dissecting the autism-associated 16p11.2 locus identifies multiple drivers in brain phenotypes and unveils a new role for the major vault protein

**DOI:** 10.1101/2022.01.23.477432

**Authors:** Perrine F. Kretz, Christel Wagner, Charlotte Montillot, Sylvain Hugel, Ilaria Morella, Meghna Kannan, Anna Mikhaleva, Marie-Christine Fischer, Maxence Milhau, Riccardo Brambilla, Yann Herault, Alexandre Reymond, Mohammed Selloum, Stephan C. Collins, Binnaz Yalcin

## Abstract

Using mouse genetic studies, we set out to identify which of the 30 genes causes brain size and other NeuroAnatomical Phenotypes (NAPs) at the autism-associated 16p11.2 locus. We show that multiple genes mapping to this region interact to regulate brain size in contrast to previous studies, with female significantly less affected. Major Vault Protein (MVP), the main component of the vault organelle, is a highly conserved protein found in higher and lower eukaryotic cells, yet its function is not understood. Here we find MVP expression highly specific to the limbic system and *Mvp* as the top driver gene of NAPs, regulating the morphology of neurons, postnatally and specifically in male. Finally, we demonstrate that the double hemideletion *Mvp*::*Mapk3* rescues NAPs, increases phosphorylation of ERK and alters behavioral performances, suggesting that MVP and ERK share the same signalling pathway, *in vivo*. Our results highlight that sex-specific cellular and neuroanatomical mechanisms must be considered in neurological disorders such as autism. Most importantly, it provides the first evidence for the involvement of the vault organelle in the regulation of the mammalian brain size and limbic structures.

## Introduction

Autism spectrum disorders (ASDs) are a group of complex neurodevelopmental diseases characterized by restricted/repetitive behaviors and a deficit in social communication. Affected children usually express autistic behaviors after 24 months of age. ASDs are well known to be sex-biased with four males diagnosed for every female^1^. Apart from a recent study on the impact of androgens on human neural stem cells^2^, the underlying sex-specific biology remains largely unknown.

The human 16p11.2 locus is susceptible to a 600 Kb deletion (OMIM #611913) which is among the most frequent known etiologies of ASDs^3^. This 16p11.2 deletion usually arises *de novo* and partly associates with abnormal brain size and other structural defects including a decreased cortical thickness restricted to male carriers^4,5^. One of the main challenges in the field is to decipher which of the 30 protein-coding genes mapping to this region underlies the NeuroAnatomical Phenotypes (NAPs) and how they interact with each other leading to ASDs.

To evaluate the contribution of each gene within the 16p11.2 locus, systematic morpholino-mediated knockdown methods in the zebrafish model identified *KCTD13* (potassium channel tetramerization domain containing 13) as a major driver of brain size^6-8^. However, recent studies in mice did not confirm this association^9,10^, questioning the relevance of the zebrafish model in this particular syndrome.

Intriguingly, *MVP*, which encodes the 100 kDa major vault protein in the interval, was found to modify NAPs via *cis* genetic epistasis with *KCTD13*^*9,11*^. Described in 1986 by Leonard H Rome and named after its arched shape reminiscent of the ceilings of cathedrals, the vault is a highly abundant and conserved organelle present in many eukaryotic cells^12^. The major vault protein is the main constituent of the vault organelle and is sufficient to give the vault its hollow barrel-like shape^13^. Furthermore, MVP is not visible as free monomers at any given time in cells but instead appears only as vault particles^14^. Indeed, Tanaka *et al*. have resolved the structure of the rat vault at 3.5 Å resolution as a particle made of 78 identical MVP^15^. This makes the mammalian vault the biggest ribonucleoprotein organelle in cells, three times the size of a ribosome, that can potentially be used as a drug delivery system^16^.

Despite its discovery more than 30 years ago, the function of the vault organelle remains elusive. *In vivo*, MVP/vault is expressed in the cytoplasm, neurites and growth cone of rat cortical neurons^17^, and associated with cortical plasticity in a monocular deprivation mouse model^18^. *In vitro*, the vault organelle binds to microtubules^19^ and MVP interacts with ERK through epidermal growth factor signaling^20,21^. Interestingly, *MAPK3* (mitogen-activated protein kinase 3) encoding the extracellular-signal related kinase 1 or ERK1, maps to the 16p11.2 locus.

Here we set out to assess, in an unbiased and systematic manner, the neuroanatomical implication of each individual gene of the 16p11.2 interval using mouse mutants to inform the genetic architecture of this autism-associated locus. We find that the major vault protein (*Mvp*) gene is one of the top drivers of neuroanatomical phenotypes, regulates neuronal size, and interacts with *Mapk3* at the locus.

## Results

### Mouse genetic studies unravel the implications of multiple drivers in NeuroAnatomical Phenotypes at the 16p11.2 locus

To identify which gene(s) regulate(s) mammalian brain architecture at the *de novo* 16p11.2 deletion locus, we aim to assess NeuroAnatomical Phenotypes (named NAPs by us^22^), independently in male and female heterozygous (het) knockout (KO) mice, for each of the 30 protein-coding genes in the interval.

We developed, or acquired through collaboration, 236 adult mutant and 204 colony-matched WT mice, representing 20 unique genes (highlighted in red in **Fig. 1A**), each studied with an average of four biological replicates. For the remaining 10 genes, the germline transmission of the mutation failed despite multiple attempts or no mouse model was available during the course of the study. For one gene of interest (*Kctd13*), multiple allelic strategies were used. To ensure high comparability between the results, mouse mutants assessed in this study were all processed on identical genetic background (C57BL/6) and at same age using the same pipelines. A detailed description of study samples and allelic constructions is provided in **Supplementary Methods** and **Extended Table 1**.

**Figure 1.**
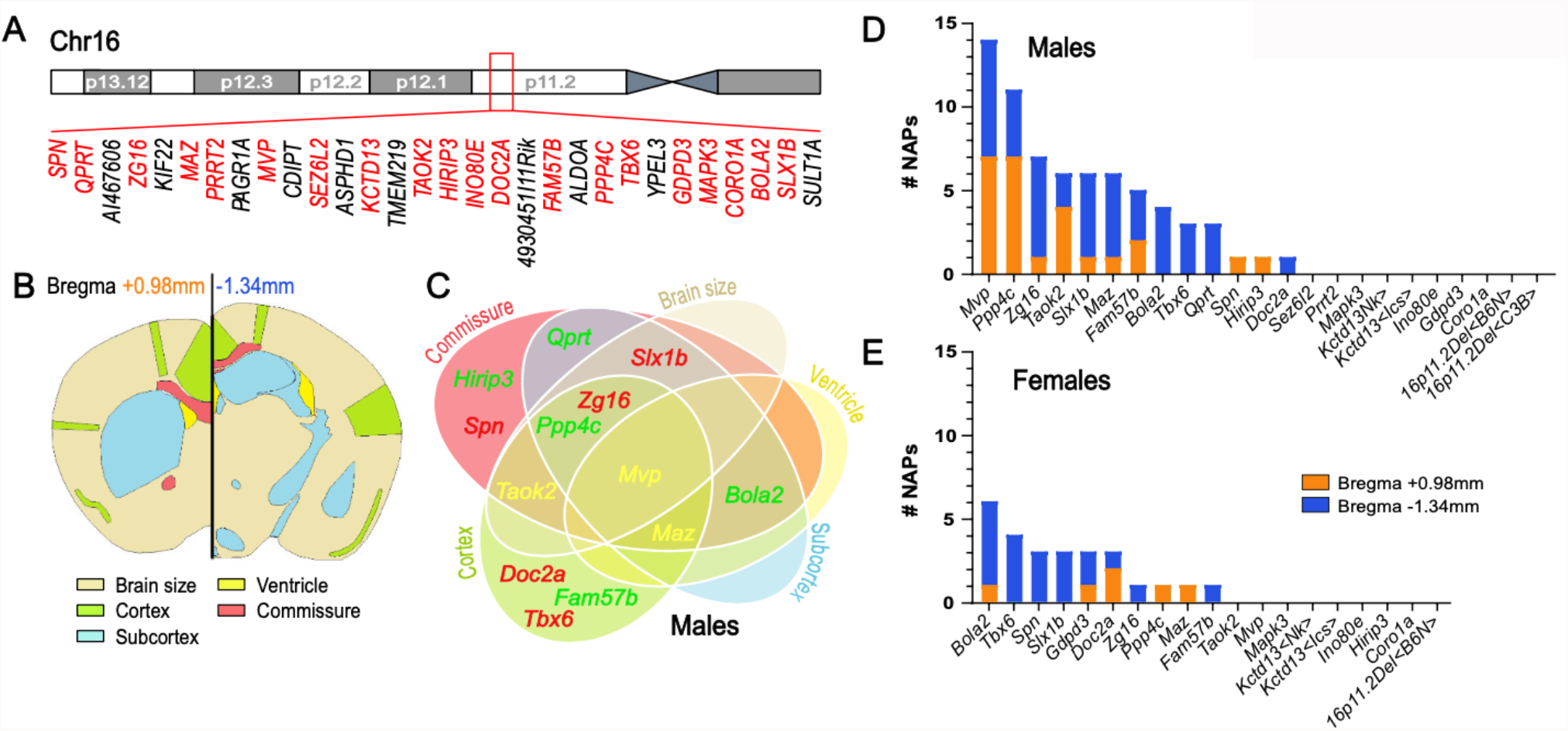
Mouse neuroanatomical studies for the identification of NAP genes at the 16p11.2 autism-associated locus. (**A**) Schematic representation of the human 16p11.2 region showing gene content and order with genes that underwent mouse neuroanatomical studies in red. (**B**) 67 brain parameters (listed in **Extended Table 2**) are grouped into five categories (brain size, ventricle, cortex, commissure and subcortex) on two coronal sections at Bregma +0.98mm and Bregma -1.34mm. (**C**) Venn diagram illustrating NeuroAnatomical Phenotype (NAP) genes (genes whose mutations yield neuroanatomical phenotypes) in male mice positioned on each category. Phenotypic directionality is color-coded. Green font corresponds to reduction in structure size, red to increase and yellow to both. Bar plots showing the number of NAPs per section (Bregma +0.98mm in orange and Bregma -1.34mm in blue) for each gene assessed for male (**D**) and female (**E**), respectively. Genes are listed on the x-axis and sorted according to the number of NAPs.

Using a highly robust approach for the assessment of 67 neuroanatomical parameters (**Extended Fig. 1**), described in details elsewhere^23^, we systematically quantified the same two coronal brain sections (Bregma +0.98 mm and Bregma −1.34 mm), and collected neuroanatomical measurements blind to the genotype (**Dataset 1**). These parameters were grouped into five main categories: brain size, commissure, ventricle, cortex and subcortex (**Fig. 1B** and **Extended Table 2**). After multiple quality control steps and critical evaluation of each phenotype, gene association was carried out within our internal database using a standardized statistical pipeline (**Supplementary Methods**). Heat maps of the assessed genes comprising percentage change relative to WTs and p-value are provided in **Dataset 2, Extended Figure 2** for male and **Extended Figure 3** for female.

We identified 13 genes associated with NAPs (hereafter named as NAP genes), when using a relaxed significance threshold of 0.05, associated with defects in commissure (10 genes), cortex (8 genes), subcortical structures (7 genes), brain size (5 genes) and ventricle (3 genes) in male (**Fig. 1C**). Eight genes (*Bola2, Qprt, Maz, Mvp, Ppp4c, Slx1b, Taok2* and *Zg16*) gave significant results affecting two or more categories. *Mvp* was the only gene affecting all five main categories. The remaining five genes (*Doc2a, Fam57b, Hirip3, Spn* and *Tbx6*) presented specific phenotypes in one brain category. In the 13 NAP genes, 38.5% decreased the size of the affected brain structures, while 38.5% increased their sizes and 23% had bidirectional effects (**Fig. 1C** and **Dataset 2**). When using a stringent significance threshold of 0.0001, *Mvp* and *Tbx6* remained NAP genes.

**Figure 1D** shows the number of NAPs for each of the 21 individually deleted allele in male. While *Mvp* still stood out as the strongest candidate gene based on the number of affected parameters (n=14), *Ppp4c, Zg16* and *Taok2*, showed ten, seven and six NAPs, respectively. These neuroanatomical phenotypes are described in details in **Supplementary Results** and a heat map provided in **Extended Figure 2**. Our top driver *Mvp* was associated with small brain size (−9%, *p*=0.016) concomitant with small brain nuclei associated to the limbic system such as the cingulate gyrus (−13%, *p*=0.0016), the somatosensory cortex (−12%, *p*=0.0025) and the hippocampus (−20%, *p*=0.023). Parameters pertaining to brain commissures were also reduced in size, for example the soma of the corpus callosum was reduced by -22% (*p*=0.044). By contrast, the size of the ventricles was enlarged by 61% (*p*=0.045) (**Extended Fig. 4A**).

Next, sex differences were assessed to determine the impact of each mutation on the female brain. Overall, female mice displayed reduced number of NAP genes (10 as opposed to 13 in male) and for each of the nine genes in common between male and female, there were less affected brain parameters in female (**Fig. 1E**). Directionality of the phenotypes was consistent between sex but NAPs were significantly different with an excess of parameters at Bregma -1.34mm for female (*p*=0.00006, Fisher test) (**Fig. 1D-E**). *Gdpd3* was a female-only NAP gene, *Bola2* showed more severe anomalies in female and *Mvp* no anomalies in female (**Extended Fig. 4B**). Female NAPs are described in details in **Supplementary Results** and **Extended Figure 3**.

Four genes (*Coro1a, Ino80e, Mapk3* and *Kctd13*) were categorized as non-NAP, both in male and female (**Extended Fig. 4C**). Considering the discrepancies in the literature^9-11^, *Kctd13* was assessed twice using two independent allelic constructions (**Supplementary Methods**), which gave identical results. We also studied the homozygous (hom) *Kctd13* mice and found a reduction in the size of the hippocampus by 10% (*p*=0.015) for male (**Extended Fig. 2** and **Extended Fig. 4D-E**) and 6% (*p*=0.014) for female (**Extended Fig. 3**). This reinforces the existing link between *Kctd13* and hippocampal biology^9,10^. Finally, to assess the genetic interactions between the 30 protein-coding genes of the interval, we studied brain anatomy in a previously published 16p11.2^+/Del^ mice^24^, in male and female (**Supplementary Methods**) but found no NAPs (**Fig. 1D-E**).

To summarize, our neuroanatomical studies demonstrate the implication of multiple major genes that drive brain size and other NAPs at the 16p11.2 locus with the major vault protein, *Mvp*, gene being one of the top drivers. Our work also indicates profound sex differences with more NAPs in male.

### The major vault protein is expressed in the limbic system and is implicated in neuronal morphology

MVP is important for the regulation of brain anatomy as evidenced by the large number of the associated NAPs when hemideleted in the mouse (**Extended Fig. 4A**). However, very little is known about how MVP could be implicated in mammalian brain biology.

To gain a comprehensive understanding of MVP/vault expression in the brain, we established its distribution at multiple scales, independently in male and female WT mice (**Supplementary Methods**). MVP transcripts, assessed at several developmental stages from embryonic day 16.5 (E16.5) to 30 weeks of age, were constant. Expression was higher in the cerebellum and peripheral tissues than in the cortex and hippocampus (**Fig. 2A** and **Extended Fig. 5A-D**). The neuroanatomical distribution of the vault organelle was then quantified by immunofluorescence using an anti-vault antibody, throughout the entire brain on consecutive histological sections. MVP/vault signal was refined to the granular layer and the arbor vitae of the cerebellum (**Fig. 2B**), the oriens layer of the CA3 region of the ventral hippocampus (**Fig. 2C**) and the deep layers of the cingulate gyrus (**Fig. 2D**). We noticed MVP/vault signal pertained to specific nuclei of the limbic system, for example the triangular septal nucleus, the zona incerta, the solitary nucleus, the vagal nucleus, the paraventricular hypothalamic nuclei and the dorsal medial thalamic nucleus (**Extended Table 3** and **Extended Fig. 5E-N**). At the subcellular level, MVP/vault signal was limited to the cytoplasm of neurons (**Fig. 2E** and **Extended Fig. 5O-Q**). It was noteworthy that the patterns observed were consistent across sectioning planes (coronal and sagittal) and replicates (**Extended Table 4**) and that no sex differences were detected in the pattern of MVP/vault expression.

**Figure 2.**
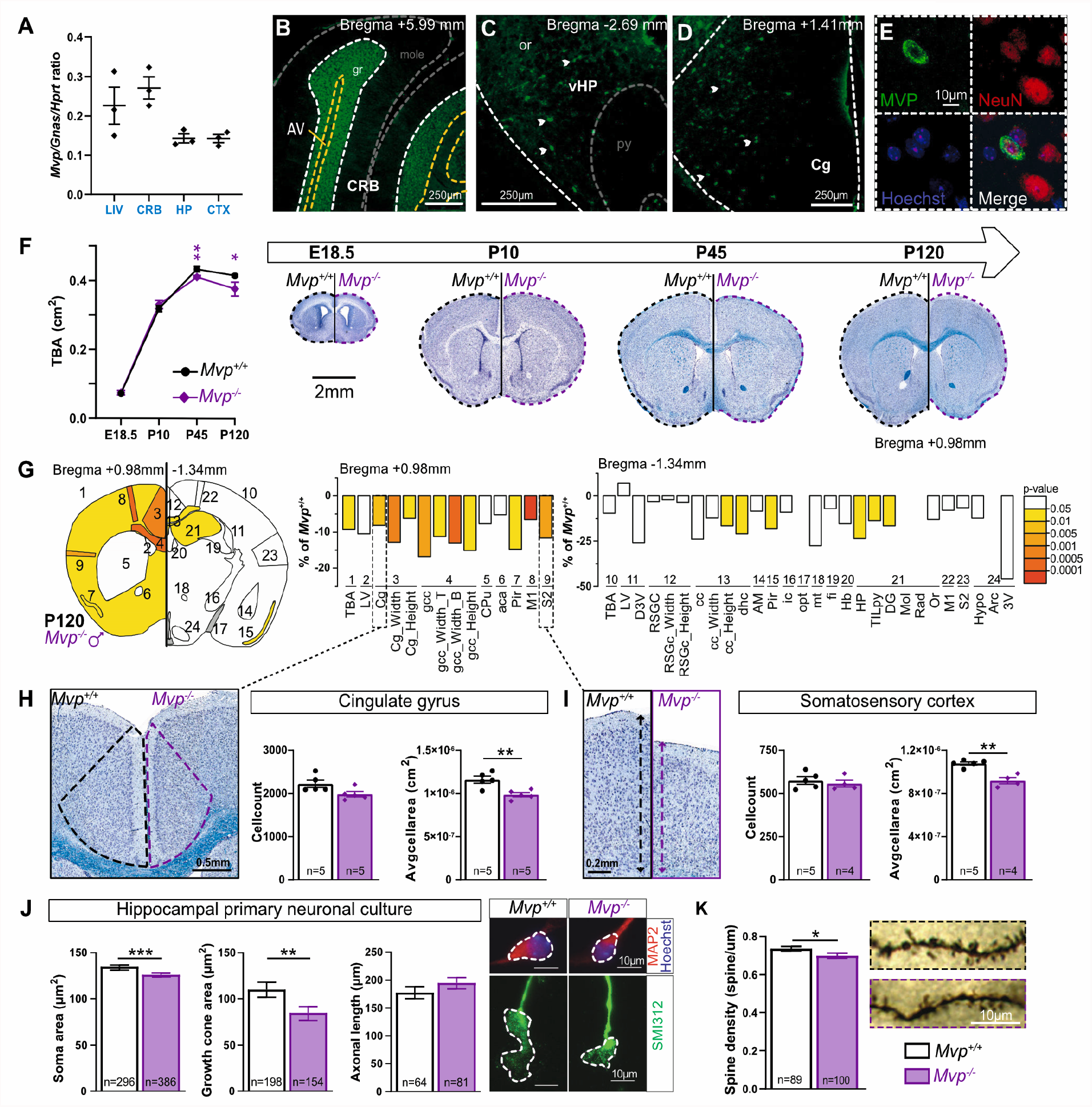
Involvement of the major vault protein (MVP) gene in male brain morphology. (**A**) *Mvp* qRT-PCR expression in liver (LIV), cerebellum (CRB), hippocampus (HP) and cortex (CTX). Normalization was done using *Gnas*/*Hprt* ratio. Immunofluorescence of MVP (green) in CRB (**B**), ventral HP (**C**) and deep layers of cingulate gyrus (Cg) (**D**). Arrow heads point to MVP-positive cells. AV, arbor vitae; gr/mole, granular/molecular layer of CRB; or/py, oriens/pyramidal layer of HP. (**E**) Image of one single MVP-positive cell (top left), co-labeled with NeuN (red) and Hoechst (blue). For detailed description of MVP expression studies, see **Extended Tables 3-4** and **Extended Figure 5**. (**F**) *Left*. Graph showing the total brain area (TBA) measured at four time points, embryonic age 18.5 (E18.5), postnatal day 10 (P10), P45 and P120. *Right*. Half images of coronal brain sections stained with Nissl-luxol at the four time points from *Mvp*^*+/+*^ and *Mvp*^*-/-*^. (**G**) *Left*. Schematic representation of 24 brain regions assessed at Bregma +0.98mm and -1.34mm in male *Mvp*^*-/-*^ mice at P120. Colored regions indicate the presence of at least one significant parameter within the brain region at the 0.05 level. White indicates a p-value >0.05, grey shows not enough data to calculate a p-value. *Right*. Histograms of percentage change relative to *Mvp*^*+/+*^ for each of the 42 parameters (see **Extended Table 2**). (**H-I**). *Left*. Coronal sections of Cg (**H**) and somatosensory cortex S2 (**I**) from *Mvp*^*+/+*^ and *Mvp*^*-/-*^ at P120. *Right*. Cell count and average cell area measures within Cg and S2. (**J**) *Left*. Graphs showing measures of soma area, growth cone area and axonal length from hippocampal neurons from *Mvp*^*+/+*^ and *Mvp*^*-/-*^. *Right*. Neurons stained with MAP2 (red) and SMI312 (green). (**K**) *Left*. Spine density from dendrites of pyramidal neurons in the cortex from *Mvp*^*+/+*^ and *Mvp*^*-/-*^. *Right*. Representative Golgi images. Plots are shown as mean ± SEM (except **G**). One-way ANOVA with Tukey *post-hoc* (**A**), two-tailed Student *t* test equal variance (**F-K**). * *p*<0.05 * * *p*<0.01 * * * *p*<0.001.

The MVP mouse model used in this study was validated as loss-of-function (LoF) of *Mvp*, assessed at the transcript and protein levels (**Extended Fig. 6A-D** and **Supplementary Methods**). We also verified that the expression of neighbouring genes was unaffected (**Extended Fig. 6E**). Among 617 successfully genotyped animals, we observed Mendelian ratio inheritance, indicating that the loss of *Mvp* has no effect on survival (**Extended Fig. 6F**).

To discriminate primary microcephaly from acquired microcephaly, we studied brain size in *Mvp*^*-/-*^ mice across four time points: embryonic day 18.5 (E18.5), postnatal day 10 (P10), P45 and P120, using an adaptation of the procedure described in the previous section (**Extended Table 5**). The total brain area measurement in male *Mvp*^*-/-*^ was normal at E18.5 and P10 but smaller at P45 (−7%, *p*=0.0061) and P120 (−9%, *p*=0.046), suggesting that the microcephaly is acquired between P10 and P45 (**Fig. 2F**). At P120, in addition to the total brain area being decreased, 16 parameters were significantly smaller including the area of the cingulate gyrus (−8%, *p*=0.007), the thickness of the somatosensory cortex (−12%, *p*=0.0016) and the area of the hippocampus (−24%, *p*=0.038) (**Fig. 2G**). In female *Mvp*^*-/-*^, no change was detected at any given time (**Extended Fig. 7A-B**). Number and effect size of NAPs being similar between *Mvp*^*-/-*^ and *Mvp*^*+/-*^ male (17 NAPs versus 14, respectively), we performed subsequent studies on *Mvp*^*-/-*^ only. Given the changes of cortical thickness, we first asked whether alterations in one specific cortical layer may contribute to this phenotype but found none (**Extended Fig. 8A-D**). NAPs in male *Mvp*^*-/-*^ were confirmed on sagittal planes using a previously described procedure^25^ (**Extended Fig. 8E** and **Extended Table 6**). To determine why brain regions were smaller in *Mvp*^*-/-*^ mice, we took advantage of the high resolution of our approach and developed a suite of automated tools to count the number of cells and calculate the average cell size within each brain region (**Supplementary Methods**). No significant change was detected in the number of cells in male *Mvp*^*-/-*^, however cells were significantly smaller in size in all affected brain regions including the cingulate gyrus (−15%, *p*=0.0053) (**Fig. 2H**) and somatosensory cortex (−15%, *p*=0.001) (**Fig. 2I**). Cell size was normal in unaffected brain region such as the retrosplenial cortex (**Extended Fig. 8F-M**). The same set of studies were conducted in female *Mvp*^*-/-*^ but neither defects in cell count nor size were found (**Extended Fig. 7C-H**).

To further characterise the cellular phenotype, we conducted hippocampal neuronal cultures, independently in male and female (**Supplementary Methods**). Consistently, neurons derived from male *Mvp*^*-/-*^ showed a reduction of the soma size by 6% (*p*=0.0003). The growth cones were smaller by 23% (*p*=0.0081) (**Fig. 2J**) and no differences were detected for the axonal length (**Fig. 2J**). Neurons derived from female *Mvp*^*-/-*^ showed no morphological defects (**Extended Fig. 7I-J**).

Finally, to test if the anatomical changes relate to neural connectivity defects in male *Mvp*^*-/-*^, we measured miniature excitatory postsynaptic currents (mEPSCs) in pyramidal neurons of the anterior cingulate gyrus (**Supplementary Methods**). Interestingly, the mEPSCs amplitude was smaller (*p*=0.023) (**Extended Fig. 9A**) and the density of postsynaptic dendritic spines was reduced (−5%, *p*=0.02) (**Fig. 2K** and **Dataset 3**), indicating functional and morphological changes in synaptic connections. Neuronal ultrastructure was examined but no anomalies were seen in the cingulate gyrus (**Extended Fig. 9C-D**). To identify potential biological pathways that might explain NAPs, we carried out transcriptomic analyses in the cingulate gyrus, and although the complete loss of *Mvp* was confirmed, no differentially expressed genes were found (**Extended Fig. 9E**).

All together, these findings show that *Mvp* is not essential for survival but has a highly specific pattern of expression in the limbic system and is implicated in the regulation of neuronal size and dendritic spines, postnatally.

### The double hemideletion of *Mvp* and *Mapk3* rescues neuroanatomical phenotypes and alters behavioral performances in mice

MVP-mediated regulation of ERK signalling has been demonstrated in two previous studies^20,21^. To explore the relevance of this interaction, we generated double KOs of *Mvp* and *Mapk3* (**Supplementary Methods**).

We first examined ERK activity by measuring phospho-ERK in the cortex of *Mvp*^*+/+*^*::Mapk3*^*+/+*^, *Mvp*^*+/+*^*::Mapk3*^*+/-*^, *Mvp*^*+/-*^*::Mapk3*^*+/+*^ and *Mvp*^*+/-*^*::Mapk3*^*+/-*^ mice. Previous work indicated that *Mapk3*^*-/-*^ showed an upregulation of ERK activity^26^. Here she show that *Mvp*^*+/+*^*::Mapk3*^*+/-*^ exhibited an increase of global ERK phosphorylation also (**Fig. 3A**), upon hemiablation of the gene *Mapk3* confirmed by a 50% reduction of ERK1 protein levels (**Extended Fig. 10A-B**), while female were unaffected (**Fig. 3B**). Our quantification of ERK activity revealed a three-fold increase of phospho-ERK in the cortex of *Mvp*^*+/-*^ *::Mapk3*^*+/+*^, indicating that MVP could be an inhibitor of ERK signaling. Accordingly, ERK activity was reduced by more than a third in the double hets *Mvp*^*+/-*^*::Mapk3*^*+/-*^. These preliminary findings provide evidence of MVP-mediated regulation of ERK signalling, *in vivo*.

**Figure 3.**
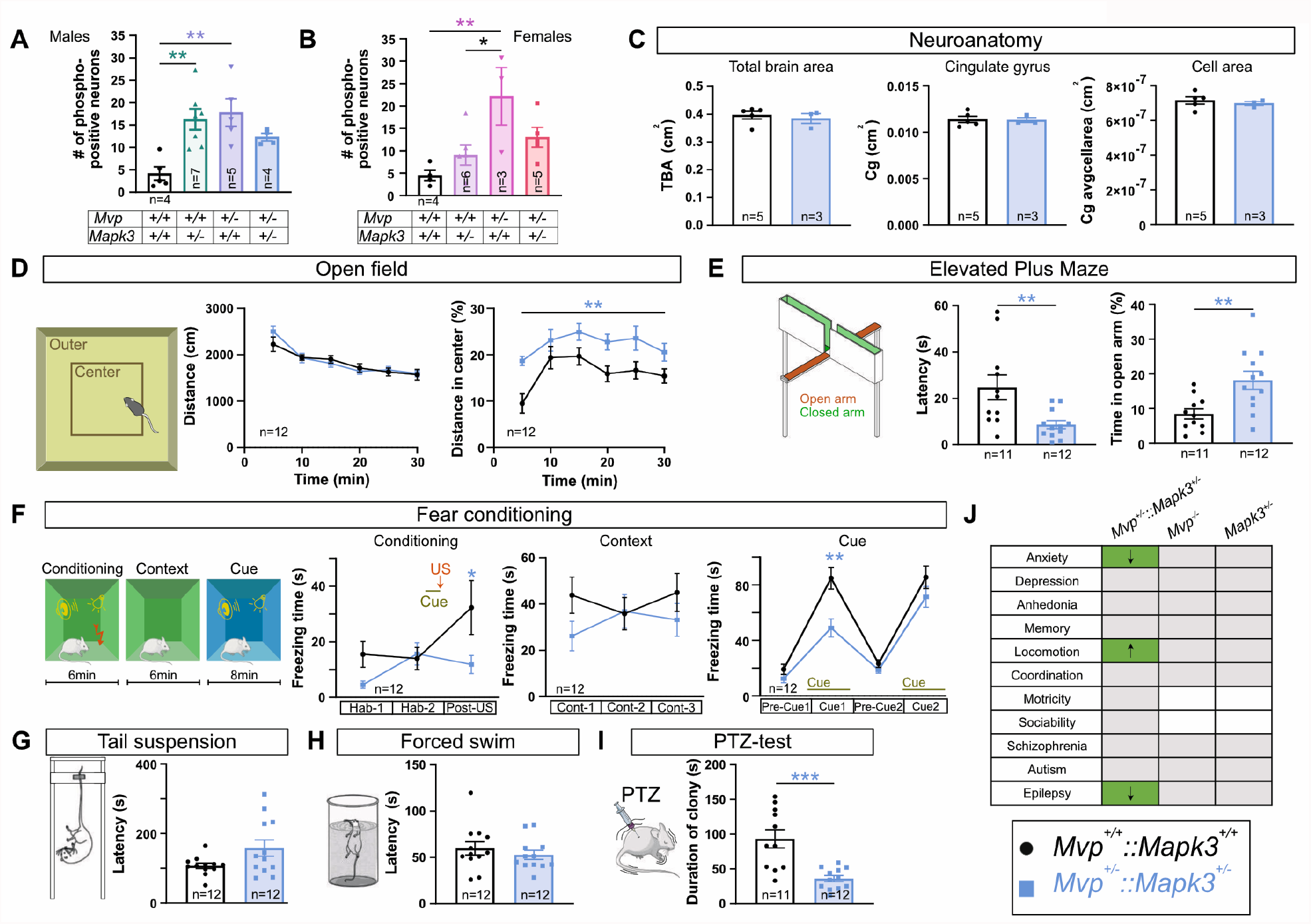
Analysis of double heterozygous hemideletion of *Mvp* and *Mapk3* genes. (**A-B**) Number of phospho-ERK positive neurons quantified through the prefrontal cortex by immunofluorescence in male (**A**) and female (**B**). (**C**) Selection of three histological parameters in *Mvp*^*+/-*^*::Mapk3*^*+/-*^ mice: total brain area (TBA), cingulate gyrus (Cg) area and average Cg cell area (see **Dataset 1** and **Dataset 3** for neuroanatomical and cellular raw data, respectively). Selection of behavioral paradigms among 17 tested in 12 *Mvp*^*+/+*^*::Mapk3*^*+/+*^ and 12 *Mvp*^*+/-*^*::Mapk3*^*+/-*^ mice, from 11 to 25 weeks of age: open field (**D**), elevated plus maze (**E**), fear conditioning (**F**), tail suspension (**G**), forced swim (**H**) and PTZ-test (**I**). (**J**) Summary table of core functions assessed in various cohorts. Green indicates significance, grey no difference and white no data. Arrows indicate directionality of effect. A comprehensive description of behavioral tests, raw datasets and additional findings are provided in **Supplementary Methods, Datasets 4-6** and **Extended Figures 10-12**, respectively. Mean and standard error of the mean are shown in the graphs and sample size indicated within each graph. Two-tailed Student *t* test equal variance (**C, E, G**-**I**), one-way ANOVA with Tukey post hoc (**A**-**B**) and two-way ANOVA with Sidak post hoc (**D, F**). * *p*<0.05 * * *p*<0.01 * * * *p*<0.001.

Next, we produced additional cohorts of double KOs of *Mvp* and *Mapk3* to assess NAPs using our standard procedures for histomorphology^27^. We found no differences in *Mvp*^*+/-*^*::Mapk3*^*+/-*^ when compared to *Mvp*^*+/+*^*::Mapk3*^*+/+*^, indicating that the decreased expression of *Mapk3* rescues the impact of *Mvp*-deficiency on brain size and anatomy (**Fig. 3C**).

Furthermore, we ran a range of sixteen behavioral tests evaluating eleven core behaviors (anxiety, depression, anhedonia, memory, locomotion, coordination, motricity, sociability, schizophrenia, autism and epilepsy) in double het and single-gene male adult KOs. The behavioral pipeline is described in details in the **Supplementary Methods** and the raw data available in **Datasets 4-6**. The double hets showed normal body weight (**Extended Fig. 10C**) as well as normal motricity, coordination, reward-seeking behaviors, working and short-term memory, associative learning, circadian activity, and no signs of autism and schizophrenic-like behaviors (**Extended Fig. 10D-M**). While the open-field test indicated no increase in traveled distance suggesting normal basic activity, it showed increased time spent in the center of the apparatus and decreased latency to enter the center (**Fig. 3D**), suggesting a resistance to anxiogenic behavior. The elevated plus maze confirmed the resistance to anxiety, shown by more time spent in the open arm of the device (**Fig. 3E**). Fear conditioning in contextual and cued testing showed decreased freezing (**Fig. 3F**), indicating improved perception skills. The tail suspension test showed a trend to increased latency to immobility (**Fig. 3G**), while the forced swim did not show any apparent differences (**Fig. 3H**). Finally, the duration of clonic seizures was dramatically reduced in the pentylenetetrazol (PTZ)-induced seizure paradigm (**Fig. 3I**), supporting the idea of a resistance to epilepsy also. Additionally, we studied the same series of eleven core functions and assessed the phenotypes of single-gene mutation of *Mvp* and *Mapk3*. We found no behavioral anomalies in *Mvp*^*-/-*^ and *Mapk3*^*+/-*^ (**Extended Fig. 11** and **Extended Fig. 12**, respectively), suggesting that the het deletion of *Mvp* or *Mapk3* is alone not sufficient to potentiate the enhanced behaviors seen in the double het KOs. A summary of behavioral findings is provided in **Figure 3J**.

Taken together, these results confirm the *in vivo* interaction between MVP and ERK1, indicate neuroanatomical correction of *Mvp*-deficiency by reducing expression of *Mapk3*, and reveal the benefit of a double hemideletion of *Mvp* and *Mapk3* in the neurobiology of fearfulness and epilepsy.

## Discussion

Our findings are important in at least three respects. First, we found multiple drivers of neuroanatomical phenotypes with opposite effect within the 16p11.2 interval that undergo complex additive and synergistic epistasis as recently reported in this region^8,28^ and other loci^29^. Among the 20 syntenic genes assessed, two were associated with decreased (−9% for *Mvp* and -10% *Ppp4c*) and three with increased (+7% for *Slx1b*, +6% for *Taok2* and +7% for *Zg16*) size of the brain. No neuroanatomical changes were detected in the double hemiablation of *Mvp*::*Mapk3* nor in 16p11.2^+/Del^ mice^24^, suggesting that the interactions between genes at the locus can hide neuroanatomical phenotypes driven by individual genes. This could explain the variable MRI findings in patients harboring the 16p11.2 deletion, with one-third to two-thirds not showing brain anatomical changes^30,31^.

We did not find any brain morphological changes in *Kctd13* het mice, consistent with two recent mouse studies^9,10^. At the exception of *Taok2*, linked to enlarged brain size in a recent report^32^ and this study, NAP genes identified were new from the literature, both in mouse and human, potentiating neurodevelopmental disease gene discovery. Among non-NAP genes, findings were consistent with a report of *Prrt2* KO mice that did not find any overt abnormalities in brain size^33^.

Second, we provide the first evidence that the vault organelle modulates brain and neuronal size in key structures of the limbic system where it is expressed including the cingulate gyrus, a structure associated with the management of emotions^34^. We found no differential expression changes in the cingulate gyrus, arguing against a major transcriptional effect of the loss of *Mvp*. MVP has previously been associated with synaptic transmission in other models^18,35^. The vault organelle might also be implicated in the transport of mRNAs along neurites for translation at the synapse^17^, although the totality of mRNA species that bind to the vault organelle remains to be identified. We found a decrease in postsynaptic spine density and mEPSCs amplitude in pyramidal cingulate neurons, suggesting that the loss of *Mvp* disrupts neural connections also.

Third, we provide insights into the mechanisms of sex differences at the 16p11.2 locus in a suitable model, sexing being not possible in fruit fly^28^ and zebrafish^8,11^. From an anatomical perspective, the female mouse brain is more robust while the male brain is more vulnerable when a genetic mutation is involved. This validates the “female protective effect” previously predicted in human neurodevelopmental disorders^36^. Our findings on MVP exclude expression differences between male and female as a potential cause responsible for the underlying sex differences in NAPs, favoring the hormonal hypothesis^37^.

ERK1/2 kinase signaling cascades are central in the control of neuronal plasticity and size^38^, and can be activated in response to testosterone^39^. While hyperphosphorylation of ERK1 has been previously reported in several studies of 16p11.2^+/Del^ mice, sex differences in ERK phosphorylation were overlooked^40,41^, at the exception of one recent study^42^. On this ground, pharmacological ERK inhibitors have been used to improve neurological traits in 16p11.2^+/Del^ mice^43^, and more generally in neurodevelopmental and neurodegenerative disorders^44^. In the current study, hemiablation of *Mapk3* showed increased ERK1 activation in male cortex not female, reinforcing a hormone-dependent regulation of kinase cascades. Our double heterozygous knockout strategy of *Mvp* and *Mapk3* supports the notion that the vault organelle could act as a naturally-occurring inhibitor of ERK1 signaling pathway to ameliorate behavioral responses to environmental stressors.

Finally, human and mouse MVP sharing approximately 90% of their amino acid residues, the vault organelle likely plays a similar role in the human brain. Human MVP alone has not been implicated in shaping brain size, possibly due to a polygenic model involving other genes of the 16p11.2 locus such as *Mapk3*. Interestingly, a smaller deletion region for autism encompassing five out of the 30 genes have been narrowed down^45^ and included *MVP*. In summary, we have begun to identify what may be an underlying reason why neurodevelopmental disorders such as autism predominantly affect boys with the identification of the vault organelle and its implication in ERK1 signaling pathway.

## Acknowledgements

We are very grateful to Leonard H Rome, Nancy Lynn Kedersha and Valerie Kickhoefer for discussions on the vault organelle and kindly providing anti-MVP IgG antibodies (George and Charlie) and anti-vault antibody (N2-B15). We thank Ipek Yalcin for discussions and her guidance in isolating the anterior cingulate gyrus. For histological work, we thank Sylvie Nguyen as well as Dorinda Wright and Lindsey Proctor at HistologiX Ltd. We thank the students involved in the study (Jules Roussey, Emeline Aguilar, Paula Hahn, Durna Kumruoglu, David Gualberto, Jonathan Delevoye, Sarah Arthur, Rebecca Balz and Léo Gagliardi). We thank Renee Araiza, Taylor J Lunceford and Kevin C. Kent Lloyd for the shipping of the *Hirip3* mouse. We thank Gilles Pagès (University of Nice), Virginia E. Papaioannou (University of Columbia), Yoshimi Takai (Kobe University), and Nicholas Katsanis and Christelle Golzio (Duke University Medical Center) for kindly providing the *Mapk3, Tbx6, Doc2a* and *Kcdt13* mice, respectively. We thank Jean Pieters (University of Basel) and Tsutomu Fukuwatari (Kyoto University) for kindly providing brain samples for *Coro1a* and *Qprt*, respectively. We thank members of the International Mouse Phenotyping Consortium (IMPC) including the Italian National Research Council (Roma), the Mouse Clinical Institute (Illkirch), the Mouse Biology Program (MBP) at UC Davis (California), the Texas A&M Institute for Genomic Medicine (TIGM) and the Wellcome Sanger Institute (Hinxton). We thank members of core facilities at the Institute of Genetics and Molecular and Cellular Biology (Illkirch) and the Mouse Clinical Institute (Illkirch) involved in the study including the team of Sylvie Jacquot for genotyping, Nadia Messadeq for electron microscopy, Dalila Ali-Hadji for animal care, Hamid Meziane for behavioural testing, and Christelle Thibault-Carpentier for transcriptomics. Sequencing was performed by the GenomEast platform, a member of the « France Génomique » consortium (ANR-10-INBS-0009). This study was funded by the Swiss National Science Foundation (grant 31003A_182632) to A.R., the ANR-10-INBS-07 PHENOMIN grant to YH, and by the French National Research Agency (ANR-11-PDOC-0029 and ANR-18-CE12-0009), the Gutenberg Circle, the grant ANR-10-LABX-0030-INRT, a French State fund managed by the Agence Nationale de la Recherche under the frame program Investissements d’Avenir ANR-10-IDEX-0002-02, and the Infrafrontier Research Infrastructure (TAOK2), to B.Y.

## Author contributions

B.Y. directed the study and acquired funding for the project. P.F.K., C.W., C.M., M.K., A.M., M.C.F., M.M. and S.C.C. performed/analyzed data for mouse neuroanatomy, neuronal cultures and behavior; S.H. performed/analyzed data for electrophysiology; I.M. performed/analyzed data for phospho-ERK in brain sections with supervision by R.B.; Y.H., A.R. and M.S. provided resources; S.C.C managed all aspects of the project. B.Y. wrote the paper together with P.F.K. All authors edited and approved the manuscript.

## Methods

Here we provide a summary of our main methods. For a full description, see **Supplementary Methods** in the Supplementary Information document.

### Study samples

The mouse syntenic 16p11.2 region *Sult1a-Spn* encompasses 30 protein-coding genes on chromosome 7. Among these, 20 (**Extended Table 1**) were incorporated in our analysis of neuroanatomical phenotyping explained in **Extended Figure 1**. The remaining ten were not available during the course of the study for reasons explained in **Supplementary Methods**. Genotyping primers were designed in our own animal facility (Mouse Clinical Institute, Illkirch, France). A list of sequences is provided in **Supplementary Methods**. Sexing was determined visually except for embryos and young postnatal age where we combined visual assessment with genetic testing using the SRY reactions. For validation purposes, a set of mutants was studied multiple times through the production of independent cohorts or at different ages.

### Generation of double knock-outs

We generated double-knockout lines by crossing single-gene mutants producing WT and double-het, as well as intermediate heterozygotes for each single-gene mutant. Four groups of mice were assessed consisting of *Mvp*^*+/+::*^*Mapk3*^*+/+*^, *Mvp*^*+/-::*^*Mapk3*^*+/+*^, *Mvp*^*+/+::*^*Mapk3*^*+/-*^ and *Mvp*^*+/-::*^*Mapk3*^*+/-*^.

### Animal welfare and ethical consideration

The mice were bred at the Mouse Clinical Institute under controlled light/dark cycles and were provided with food and water ad libitum. All animal procedures were carried out in accordance with the ARRIVE (Animal Research: Reporting of In Vivo Experiments) guidelines. Animal procedures were approved by the local ethics committee (Com’Eth) under the reference number 2016010717527861.

### Brain sample processing

The brains were taken from adult mice on a high-throughput phenotyping project, where each mouse was characterized by a series of standardized operating procedures^25,27^. The collection of brain samples was performed blind to the genotype. Brains from at least four mice per genotype and per gender were collected. Control brains were systematically collected within each of the mutant lines. Every aspect of the procedure was managed through a relational database using the FileMaker (FM) Pro database management system (detailed elsewhere^22^). A similar approach was used for embryos at E18.5 fixed in Bouin solution.

### Neuroanatomical studies and quality control

An overview of sample processing is shown in **Extended Figure 1**. Brains were cut at a thickness of 5μm on a sliding microtome (HM450, Microm, France) on symmetrical and stereotaxic planes to obtain coronal sections matching Bregma +0.98mm and Bregma -1.34 mm. Our precision was estimated to be no more than 30μm, anterior or posterior, to the histological section of interest. The sections were doubled-stained with 0.1% Luxol Fast Blue (Solvent Blue 38, Sigma-Aldrich) and 0.1% Cresyl violet acetate (Sigma-Aldrich), in order to label myelin and neurons, respectively. After mounting on slides, the sections were scanned at cell-level resolution using the Nanozoomer whole-slide scanner 2.0HT C9600 series (Hamamatsu Photonics, Shizuoka, Japan).

Each image was quality controlled to assess whether (i) the section is at the correct position, (ii) the section is symmetrical, (iii) the staining is of good quality, and (iv) the image is good quality. Only images that fulfilled all of the quality control checks were fully processed. These quality control steps are essential for the detection of small to moderate neuroanatomical phenotypes (NAPs) and without which the large majority of NAPs would be missed. This is explained in great details elsewhere^22^. On coronal plane, 67 brain morphological parameters of 39 area and 28 length measurements were taken for each sample (**Extended Table 2**). 42 neuroanatomical coronal measurements were taken at E18.5 (**Extended Table 5**) and 40 on sagittal plane (**Extended Table 6**). Cellular features across histological sections were measured using a semi-automated macro designed in Fiji measuring cell counts as well as averaged cell area for each cell.

### MVP expression

Several mouse cohorts underwent microdissection in order to separate various brain structures of interest at various developmental and adulthood stages. Total RNA was extracted using a phenol-chloroform technique. 1µg of total RNA was reverse-transcribed to complementary DNA using Supercript III First-Strand Synthesis Supermix (11752–050, Invitrogen) for qualitative and quantitative reverse transcription PCR (RT-PCR). For the latter, we used TaqMan assays.

### Immuno-fluorescence

Adult mice were anesthetized, intracardially perfused with PBS and fixed with 4% Paraformaldehyde (PFA) solution in PBS. Brains were harvested, post-fixed in 4% PFA for 24 hours at room temperature (RT), and then transferred in 30% sucrose solution for 24 hours at 4°C. Each brain was trimmed either for coronal or sagittal sectioning and embedded in Cryomatrix (Thermo Scientific) with a fast freeze device PrestoChill (Millestone). Brains were sectioned throughout the entire tissue blocks with a cryo-microtome (K400 station with dry ice on HM450 microtome, Microm). Immunostainings for MVP were realized without detergent using floating sections. The permeabilisation step was done by incubating brain sections in methanol at -20°C for 10 minutes. Sections were rinsed in PBS, blocked in a solution with 10% Normal Donkey Serum (NDS) and 1% Bovine Serum Albumine (BSA) in PBS and incubated with rabbit polyclonal vault antibodies (N2-B15, provided by Leonard Rome, dilution 1:1000) and mouse anti-NeuN antibodies (MAB377, Millipore, dilution 1:1000) in 0.1%NDS/PBS at 4 °C overnight under agitation. After washing with PBS, the sections were incubated for three hours RT with fluorescence-conjugated secondary antibodies coupled to anti-rabbit-Alexa-488 and anti-mouse-Alexa-647 (1:1000, Thermo Fischer Scientific). Images were acquired using confocal microscope (TCS SP5, Leica) at 20× or 80x magnification and analyzed using ImageJ.

Phospho-ERK immunohistochemistry was carried out on floating sections. After quenching with 3% H_2_O_2_, 10% methanol for 15 minutes, sections were rinsed in TBS and incubated for 1 hour in blocking solution (5% normal goat serum, 0.1% Triton X-100). Sections were then incubated overnight at 4°C with anti-phospho-p44/42 MAP kinase (Thr202/Tyr204) (1:1000, Cell Signalling Technology, Danvers, MA). On the following day, a biotinylated goat anti-rabbit secondary antibody (1:200, Vector Labs) was applied to the sections for 2 hours at room temperature. Detection of the bound antibodies was carried out using a standard peroxidase-based method (ABC-kit, Vectastain, Vector Labs), followed by incubation with DAB and H_2_O_2_ solution. The sections were subsequently dehydrated using increasing concentrations of ethanol and mounted with DPX. Images were acquired from the prefrontal cortex using a bright field microscope (Leica DMI6000B Macro/Micro imaging system) under a 40X magnification and analysed with ImageJ.

### Western blot

Cortex and liver from adult mice were dissected and homogenized with 300 μL of lysis buffer containing 1× RIPA buffer (ThermoFischer), phenylmethylsulfonyl fluoride 1%, sodium orthovanadate 1%, and protease inhibitor 1% in tubes containing ceramic beads (Precellys Lysing Kit). The tubes were incubated for 30 minutes at 4 °C and centrifuged for 20 minutes at 17,000 × g at 4 °C, and the supernatant was isolated for Western blotting. 60 μg of protein was separated on 10% SDS/PAGE (Mini-PROTEAN TGX Gels,12%,10-well, 4561043, BIO-RAD) and transferred onto nitrocellulose membrane (#1620115, BIO-RAD). Membranes were blocked with 1% BSA diluted in tris buffered saline with Tween 20 (50 mM Tris, 150 mM NaCl, 0.05% Tween 20) and probed using anti-MVP (named “George”, provided by Leonard H Rome) overnight at 4 °C. Antibody–protein interactions were revealed using chemiluminesence (RPN2108, GE Healthcare), and relative protein expression was quantified using ImageJ.

### Primary neuronal culture

The hippocampus was harvested from embryos at day 18.5 (E18.5) for cell dissociation. Between 20,000 and 30,000 cells were plated on Poly-L lysine coated coverslips in 24-well plates and incubated at 37°C with 5% CO_2_. After 4 days of *in vitro* culture (DIV4), the cells were fixed with 4% PFA in 6% sucrose for 15 minutes, and then stored in EtOH at 4°C until use. Fixed cells were incubated O/N at 4°C with primary antibodies (rabbit polyclonal anti-MAP2 (AB5622, Millipore) and mouse monoclonal antibody against pan-axonal neurofilament (SMI-312R, Covance) both diluted at 1:1000 in saturation solution (0.2% Triton, 1% BSA, 10% Normal Donkey Serum (S2170, Dutscher) in Tris-Buffered Saline). Donkey anti-rabbit coupled to Alexa647 (ab150075, Abcam) and donkey anti-mouse coupled to Alexa488 (ab150105, Abcam) were used as secondary antibodies in saturation solution without Triton. Nuclei were stained with Hoechst 3342 (1:10,000 dilution, Sigma-Aldrich). Images were acquired using a regular epifluorescence microscope 100x (Leica) at magnification of 0.55x and analysed using ImageJ.

### Golgi staining

GolgiCox staining was performed using the FD Rapid GolgiStain Kit (FD NeuroTechnologies, Ellicott City, MD) on entire fresh brains, and processed as indicated by the manufacturer. After impregnation, brains were embedded in 3% low-melting agarose and cut with a vibratome (VT1200S, Leica). 100µm thick sections were mounted on gelatine-coated slides and allow to dry for two days before staining. Images were acquired with the Hamamatsu slide scanner at 40x resolution with multi-layer mode. All the measurements were taken by the same operator, blind to the genotype using the Hamamatsu NDPviewer2.0 software.

### Behavioral experiments

Eleven core phenotypes were tested: i) anxiety in the open-field and elevated plus maze paradigms, ii) learning and memory in the Y-maze, the novel object recognition test, the three-chambers paradigm and the cued and contextual fear conditioning test, iii) locomotion in the circadian activity test, iv) epilepsy in the pentylenetetrazol (PTZ)-induced seizure paradigm, v) coordination in the rotarod performance test, vi) motricity in the grip strength test, vii) schizophrenia in the prepulse inhibition and startle reflex test, viii) autism in the social behavior and marble burying paradigms, ix) depression in the forced swim and tail suspension tests, x) anhedonia in the sucrose preference test, and xi) sociability using the social recognition test. Each test is described in details in **Supplementary Methods**.

### Statistics

Statistical analyses were performed with GraphPad Prism 8.0.2, using two-tailed Student’s *t*-tests of equal variances (except for **Extended Figure 9A**, *t*-tests of unequal variances), one-way or two-way ANOVA followed by post doc tests. Tests performed are indicated in the figure legends. Results are reported as box plots with individual data points overlaid as mean ± sample standard error of the mean, except for **Figure 2, Extended Figure 4, Extended Figure 7B** and **Extended Figure 8E**, which show average for simplicity purposes. Figures were prepared using Affinity Designer. Significance was reported as follows: * p<0.05, * * p<0.01 and * * * p<0.001. All replicates are biological replicates. For qPCR data, delta C_T_ values were obtained by normalizing C_T_ values of *Mvp* against two references genes, *Gnas* and *Hprt*.

### Data availability

All datasets generated during the current study that support its findings are available in the Source Dataset files or the accompanying **Supplementary Information** document.

## References

1 Loomes, R., Hull, L. & Mandy, W. P. L. What Is the Male-to-Female Ratio in Autism Spectrum Disorder? A Systematic Review and Meta-Analysis. J Am Acad Child Adolesc Psychiatry 56, 466–474, doi:10.1016/j.jaac.2017.03.013 (2017).

2 Quartier, A. et al. Genes and Pathways Regulated by Androgens in Human Neural Cells, Potential Candidates for the Male Excess in Autism Spectrum Disorder. Biol Psychiatry 84, 239–252, doi:10.1016/j.biopsych.2018.01.002 (2018).

3 Weiss, L. A. et al. Association between microdeletion and microduplication at 16p11.2 and autism. N Engl J Med 358, 667–675, doi:10.1056/NEJMoa075974 (2008).

4 Maillard, A. M. et al. The 16p11.2 locus modulates brain structures common to autism, schizophrenia and obesity. Mol Psychiatry 20, 140–147, doi:10.1038/mp.2014.145 (2015).

5 Martin-Brevet, S. et al. Quantifying the Effects of 16p11.2 Copy Number Variants on Brain Structure: A Multisite Genetic-First Study. Biol Psychiatry 84, 253–264, doi:10.1016/j.biopsych.2018.02.1176 (2018).

6 Blaker-Lee, A., Gupta, S., McCammon, J. M., De Rienzo, G. & Sive, H. Zebrafish homologs of genes within 16p11.2, a genomic region associated with brain disorders, are active during brain development, and include two deletion dosage sensor genes. Dis Model Mech 5, 834–851, doi:10.1242/dmm.009944 (2012).

7 Golzio, C. & Katsanis, N. Genetic architecture of reciprocal CNVs. Curr Opin Genet Dev 23, 240–248, doi:10.1016/j.gde.2013.04.013 (2013).

8 McCammon, J. M., Blaker-Lee, A., Chen, X. & Sive, H. The 16p11.2 homologs fam57ba and doc2a generate certain brain and body phenotypes. Hum Mol Genet 26, 3699–3712, doi:10.1093/hmg/ddx255 (2017).

9 Arbogast, T. et al. Kctd13-deficient mice display short-term memory impairment and sex-dependent genetic interactions. Hum Mol Genet 28, 1474–1486, doi:10.1093/hmg/ddy436 (2019).

10 Escamilla, C. O. et al. Kctd13 deletion reduces synaptic transmission via increased RhoA. Nature 551, 227–231, doi:10.1038/nature24470 (2017).

11 Golzio, C. et al. KCTD13 is a major driver of mirrored neuroanatomical phenotypes of the 16p11.2 copy number variant. Nature 485, 363–367, doi:10.1038/nature11091 (2012).

12 Kedersha, N. L. & Rome, L. H. Isolation and characterization of a novel ribonucleoprotein particle: large structures contain a single species of small RNA. J Cell Biol 103, 699–709 (1986).

13 Stephen, A. G. et al. Assembly of vault-like particles in insect cells expressing only the major vault protein. J Biol Chem 276, 23217–23220, doi:10.1074/jbc.C100226200 (2001).

14 Mrazek, J. et al. Polyribosomes are molecular 3D nanoprinters that orchestrate the assembly of vault particles. ACS Nano 8, 11552–11559, doi:10.1021/nn504778h (2014).

15 Tanaka, H. et al. The structure of rat liver vault at 3.5 angstrom resolution. Science 323, 384–388, doi:10.1126/science.1164975 (2009).

16 Buehler, D. C. et al. Bioengineered vaults: self-assembling protein shell-lipophilic core nanoparticles for drug delivery. ACS Nano 8, 7723–7732, doi:10.1021/nn5002694 (2014).

17 Paspalas, C. D. et al. Major vault protein is expressed along the nucleus-neurite axis and associates with mRNAs in cortical neurons. Cereb Cortex 19, 1666–1677, doi:10.1093/cercor/bhn203 (2009).

18 Ip, J. P. K. et al. Major Vault Protein, a Candidate Gene in 16p11.2 Microdeletion Syndrome, Is Required for the Homeostatic Regulation of Visual Cortical Plasticity. J Neurosci 38, 3890–3900, doi:10.1523/JNEUROSCI.2034-17.2018 (2018).

19 Eichenmuller, B. et al. Vaults bind directly to microtubules via their caps and not their barrels. Cell Motil Cytoskeleton 56, 225–236, doi:10.1002/cm.10147 (2003).

20 Kim, E. et al. Crosstalk between Src and major vault protein in epidermal growth factor-dependent cell signalling. FEBS J 273, 793–804, doi:10.1111/j.1742-4658.2006.05112.x (2006).

21 Kolli, S., Zito, C. I., Mossink, M. H., Wiemer, E. A. & Bennett, A. M. The major vault protein is a novel substrate for the tyrosine phosphatase SHP-2 and scaffold protein in epidermal growth factor signaling. J Biol Chem 279, 29374–29385, doi:10.1074/jbc.M313955200 (2004).

22 Collins, S. C. et al. Large-scale neuroanatomical study uncovers 198 gene associations in mouse brain morphogenesis. Nat Commun 10, 3465, pdoi:10.1038/s41467-019-11431-2 (2019).

23 Mikhaleva, A., Kannan, M., Wagner, C. & Yalcin, B. Histomorphological Phenotyping of the Adult Mouse Brain. Curr Protoc Mouse Biol 6, 307–332, doi:10.1002/cpmo.12 (2016).

24 Arbogast, T. et al. Reciprocal Effects on Neurocognitive and Metabolic Phenotypes in Mouse Models of 16p11.2 Deletion and Duplication Syndromes. PLoS Genet 12, e1005709, doi:10.1371/journal.pgen.1005709 (2016).

25 Collins, S. C. et al. A Method for Parasagittal Sectioning for Neuroanatomical Quantification of Brain Structures in the Adult Mouse. Curr Protoc Mouse Biol 8, e48, doi:10.1002/cpmo.48 (2018).

26 Mazzucchelli, C. et al. Knockout of ERK1 MAP kinase enhances synaptic plasticity in the striatum and facilitates striatal-mediated learning and memory. Neuron 34, 807–820, doi:10.1016/s0896-6273(02)00716-x (2002).

27 Mikhaleva, A., Kannan, M., Wagner, C. & Yalcin, B. Histomorphological Phenotyping of the Adult Mouse Brain. Curr Protoc Mouse Biol 6, 307–332, doi:10.1002/cpmo.12 (2016).

28 Iyer, J. et al. Pervasive genetic interactions modulate neurodevelopmental defects of the autism-associated 16p11.2 deletion in Drosophila melanogaster. Nat Commun 9, 2548, doi:10.1038/s41467-018-04882-6 (2018).

29 Pizzo, L. et al. Rare variants in the genetic background modulate cognitive and developmental phenotypes in individuals carrying disease-associated variants. Genet Med 21, 816–825, doi:10.1038/s41436-018-0266-3 (2019).

30 D’Angelo, D. et al. Defining the Effect of the 16p11.2 Duplication on Cognition, Behavior, and Medical Comorbidities. JAMA Psychiatry 73, 20–30, doi:10.1001/jamapsychiatry.2015.2123 (2016).

31 Zufferey, F. et al. A 600 kb deletion syndrome at 16p11.2 leads to energy imbalance and neuropsychiatric disorders. J Med Genet 49, 660–668, doi:10.1136/jmedgenet-2012-101203 (2012).

32 Richter, M. et al. Altered TAOK2 activity causes autism-related neurodevelopmental and cognitive abnormalities through RhoA signaling. Mol Psychiatry 24, 1329–1350, doi:10.1038/s41380-018-0025-5 (2019).

33 Michetti, C. et al. The PRRT2 knockout mouse recapitulates the neurological diseases associated with PRRT2 mutations. Neurobiol Dis 99, 66–83, doi:10.1016/j.nbd.2016.12.018 (2017).

34 Pessoa, L. A Network Model of the Emotional Brain. Trends Cogn Sci 21, 357–371, doi:10.1016/j.tics.2017.03.002 (2017).

35 Herrmann, C., Volknandt, W., Wittich, B., Kellner, R. & Zimmermann, H. The major vault protein (MVP100) is contained in cholinergic nerve terminals of electric ray electric organ. J Biol Chem 271, 13908–13915, doi:10.1074/jbc.271.23.13908 (1996).

36 Jacquemont, S. et al. A higher mutational burden in females supports a “female protective model” in neurodevelopmental disorders. Am J Hum Genet 94, 415–425, doi:10.1016/j.ajhg.2014.02.001 (2014).

37 Baron-Cohen, S. et al. Why are autism spectrum conditions more prevalent in males? PLoS Biol 9, e1001081, doi:10.1371/journal.pbio.1001081 (2011).

38 Mazzucchelli, C. & Brambilla, R. Ras-related and MAPK signalling in neuronal plasticity and memory formation. Cell Mol Life Sci 57, 604–611, doi:10.1007/PL00000722 (2000).

39 Ransome, M. I. & Boon, W. C. Testosterone-induced adult neurosphere growth is mediated by sexually-dimorphic aromatase expression. Front Cell Neurosci 9, 253, doi:10.3389/fncel.2015.00253 (2015).

40 Pucilowska, J. et al. The 16p11.2 deletion mouse model of autism exhibits altered cortical progenitor proliferation and brain cytoarchitecture linked to the ERK MAPK pathway. J Neurosci 35, 3190–3200, doi:10.1523/JNEUROSCI.4864-13.2015 (2015).

41 Silingardi, D. et al. ERK pathway activation bidirectionally affects visual recognition memory and synaptic plasticity in the perirhinal cortex. Front Behav Neurosci 5, 84, pdoi:10.3389/fnbeh.2011.00084 (2011).

42 Grissom, N. M. et al. Male-specific deficits in natural reward learning in a mouse model of neurodevelopmental disorders. Mol Psychiatry 23, 544–555, doi:10.1038/mp.2017.184 (2018).

43 Pucilowska, J. et al. Pharmacological Inhibition of ERK Signaling Rescues Pathophysiology and Behavioral Phenotype Associated with 16p11.2 Chromosomal Deletion in Mice. J Neurosci 38, 6640–6652, doi:10.1523/JNEUROSCI.0515-17.2018 (2018).

44 Albert-Gasco, H., Ros-Bernal, F., Castillo-Gomez, E. & Olucha-Bordonau, F. E. MAP/ERK Signaling in Developing Cognitive and Emotional Function and Its Effect on Pathological and Neurodegenerative Processes. Int J Mol Sci 21, doi:10.3390/ijms21124471 (2020).

45 Crepel, A. et al. Narrowing the critical deletion region for autism spectrum disorders on 16p11.2. Am J Med Genet B Neuropsychiatr Genet 156, 243–245, doi:10.1002/ajmg.b.31163 (2011).

